# Shifting Baselines: Physiological legacies contribute to the response of reef coral to frequent heat waves

**DOI:** 10.1101/2020.04.23.056457

**Authors:** Christopher B. Wall, Contessa A. Ricci, Alexandra D. Wen, Bren E. Ledbetter, Delania E. Klinger, Laura D. Mydlarz, Ruth D. Gates, Hollie M. Putnam

**Affiliations:** University of Hawai‘i at Mānoa, Hawai‘i Institute of Marine Biology, PO Box 1346, Kāne‘ohe, HI 96744, USA; Pacific Biosciences Research Center, University of Hawai‘i at Mānoa, 1933 East West Road, Honolulu, HI 96816, USA; University of Texas at Arlington, Department of Biology, Arlington, TX 76019, USA; University of Miami, Rosenstiel School of Marine and Atmospheric Science, Miami, FL 33149, USA; University of Rhode Island, Department of Biological Sciences, Kingston, RI USA

**Keywords:** immunity, physiology, El Niño, bleaching, Symbiodiniaceae

## Abstract

1. Global climate change is altering coral reef ecosystems. Notably, marine heat waves are producing widespread coral bleaching events that are increasing in frequency, with projections for annual bleaching events on reefs worldwide by mid-century.
2. The response of corals to elevated seawater temperatures can be modulated by abiotic factors at site of origin and dominant endosymbiont type, which can result in a shift in multiple coral traits and drive physiological legacy effects that influence the trajectory of reef corals under subsequent thermal stress events. It is critical, therefore, to evaluate the potential for shifting physiological and cellular baselines driven by these factors in *in situ* bleaching (and recovery) events. Here, we use the back-to-back regional bleaching events of 2014 and 2015 in the Hawaiian Islands and subsequent recovery periods to test the hypothesis that coral multivariate trait space (here termed physiotype, sensu (Van Straalen, 2003) shift in multiple bleaching events, modulated by both environmental histories and symbiotic partnerships (Symbiodiniaceae).
3. Despite fewer degree heating weeks in the first-bleaching event relative to the second (7 *vs*. 10), bleaching severity in a dominant reef building coral on Hawaiian reefs, *Montipora capitata*, was greater (~70% *vs*. 50% bleached cover) and differences due to environmental history (reef site) were more pronounced. Melanin, an immune cytotoxic response, provided an initial defense during the first event, potentially priming antioxidant activity, which peaked in the second-bleaching event (i.e., a legacy effect). While magnitude of bleaching differed, immune response patterns were shared among corals harboring heat-sensitive and heat-tolerant Symbiodiniaceae. This supports a pattern of increased constitutive immunity in corals resulting from repeat bleaching events, with greater specialized enzymes (catalase, peroxidase, superoxide dismutase) and attenuated melanin synthesis.
4. This study demonstrates bleaching events have implications for reef corals beyond shaping their ecological assemblages. These events can change the magnitude and/or identity of response variables contributing to physiotype, thus generating physiological legacies carried over into the future. Quantifying baseline coral physiotypes and tracking their shifts will be critical to understanding and forecasting the effects of increased bleaching frequency on coral biology and ecology in the Anthropocene.

## 1. Introduction

Environmental change at local and global scales is degrading coral reef ecosystems. Notably, long-term trends in ocean warming and punctuated marine heat waves are destabilizing the symbiosis between corals and their endosymbiont algae (Symbiodiniaceae) and increasing coral bleaching events (Hughes et al., 2018a). The cumulative impacts of these episodic heat waves not only shift ecological and functional baselines (Hughes et al., 2019; McWilliam, Pratchett, Hoogenboom, & Hughes, 2020), but may also have legacy effects on coral biology, such that responses to bleaching events can be modulated by prior exposure (e.g., Brown, Dunne, Goodson, & Douglas, 2000). It is growing increasingly clear that susceptibility to bleaching in reef corals is based on the interaction of environmental histories (Brown et al., 2000; Safaie et al., 2018) and holobiont traits, including endosymbiont communities and genetics and physiology of the coral host (Barshis et al., 2013; Palmer, Bythell, & Willis, 2010; Palumbi, Barshis, Traylor-Knowles, & Bay, 2014; Sampayo, Ridgway, Bongaerts, & Hoegh-Guldberg, 2008).

Thermotolerant Symbiodiniaceae can confer bleaching resistance in some corals (Sampayo et al., 2008), and the Hawaiian coral *Montipora capitata* exhibits reduced bleaching when dominated by *Durusdinium* as opposed to *Cladocopium* Symbiodiniaceae (Cunning, Ritson-Williams, & Gates, 2016). However, corals are metaorganisms, and host properties such as constitutive immunity (Palmer, 2018a) and antioxidant capacity (Barshis et al., 2013) are also central to maintaining cellular homeostasis and preventing bleaching mortality (Palmer et al., 2010). As such, immunological processes are implicated as targets for natural selection and are integral to the future of reef-building corals in the face of climate change (Mydlarz, McGinty, & Harvell, 2010; Palmer, 2018a; Pinzón, Beach-Letendre, Weil, & Mydlarz, 2014). The nexus of these influential properties of coral stress responses – symbiont community and host immunity – and their role in repetitive natural bleaching and recovery events remains to be fully understood, especially with respect to the role of environmental history (Palumbi et al., 2014; Safaie et al., 2018). As global climate change increases the frequency and severity of coral bleaching events (Hughes et al., 2018), understanding the symbiotic, immune, and antioxidant responses of corals will be central to understanding the cumulative and latent effects of bleaching on the function of reef corals through time and enhancing our capacity to project community responses to thermal stress events.

The widely documented recent major bleaching of the Great Barrier Reef (2016-2017) reduced coral cover by >50% (Stuart-Smith, Brown, Ceccarelli, & Edgar, 2018), altered coral (Hughes et al., 2018b) and fish (Robinson, Wilson, Jennings, & Graham, 2019) assemblages, and disturbed the stock-recruitment capacity of reef corals (Hughes et al., 2019). As climate change continues to intensify, it is now critical to track shifts in the biology and function of corals through examination of multivariate trait-space, defined as ‘physiotypes’ by Van Straalen (2003; e.g., Figure 1a), the term which we use throughout this manuscript. Trait-space approaches have recently been applied to track changes in reef coral communities and their representative trait diversity following bleaching (Hughes et al., 2018b; McWilliam et al., 2020), and the generation of physiotype time series data will provide a critical match to further understand and interpret the growing collection of coral reef ecological time series datasets (De’ath, Fabricius, Sweatman, & Puotinen, 2012; Edmunds et al., 2014; Edmunds et al., 2014; McClanahan, Ateweberhan, & Omukoto, 2007). Together, these timeseries will allow us to integrate across cellular, ecological and evolutionary scales and improve our mechanistic understanding of these scale linkages. Importantly, tracking of multiple, interactive coral physiological traits provides the opportunity to reveal the potential for beneficial acclimatization or negative legacy effects that may underlie shifts in physiotype (e.g., Figure 1a) and testable mechanistic hypotheses underpinning resistance or susceptibility to repeated stress (Figure 1b).

**Figure 1.**
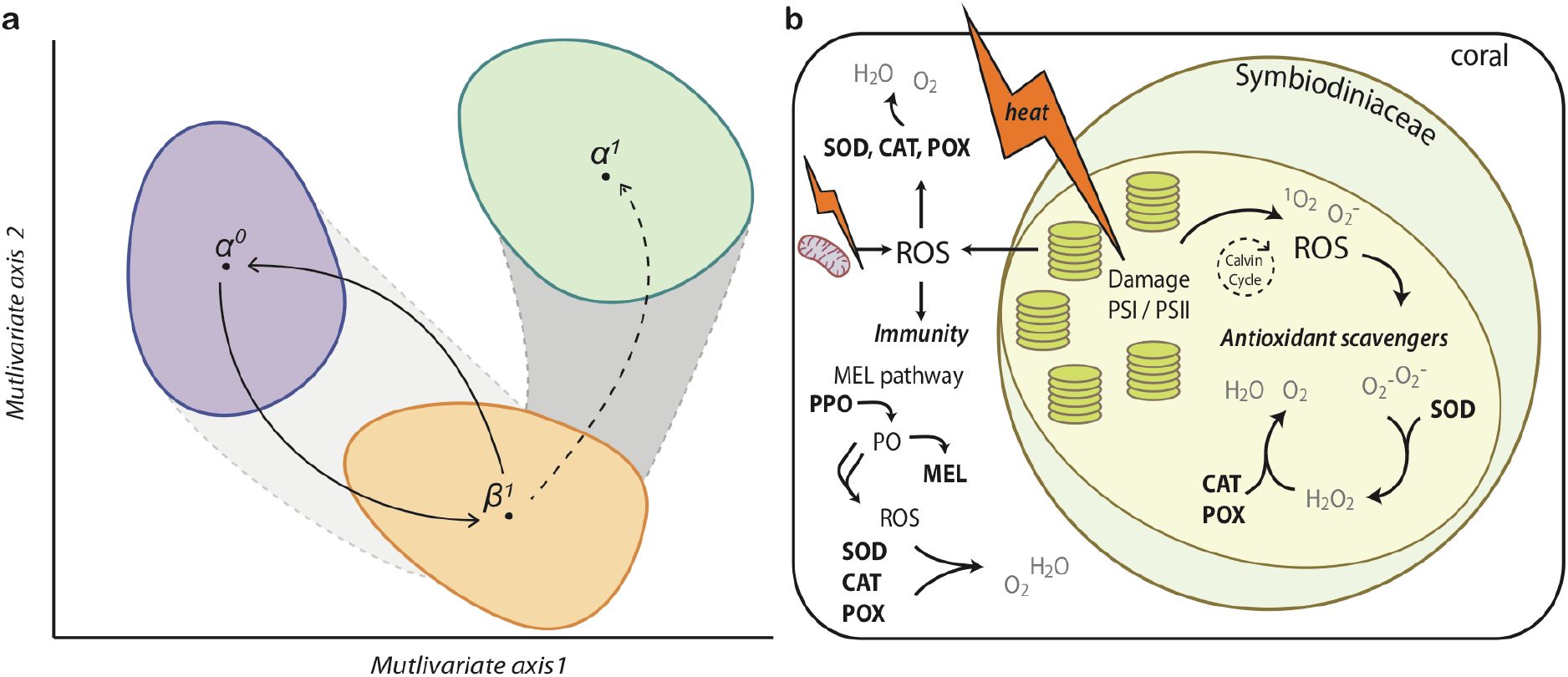
(a) Multivariate analyses identify changes in organism physiotype through space and time by accounting for the variation in multiple traits (Van Straalen, 2003). Transitions of organisms from initial (*α^0^*) to stressed states (*β^1^*), such as during bleaching in response to a heat wave, and their recovery to initial (*α^0^*) (i.e., physiological resilience) or altered states (*α^1^*). Alternatively the absence of physiotype shifts may be observed in stress resistant individuals (i.e., maintained at *α^0^*). (b) Conceptual model of melanin and antioxidant activity in coral and Symbiodiniaceae under heat stress (indicated by lightning bolts). Reactive oxygen species (ROS), oxygen singlets (^1^O_2_) and superoxide (O_2_^−^), generated by symbiont photochemical dysfunction and host mitochondria membrane damage, are neutralized by a combination of antioxidant scavenger enzymes (e.g., superoxide dismutase [SOD], catalase [CAT], and peroxidase [POX]). Host immunity via the melanin-synthesis pathway can also scavenge ROS during intermediate steps that lead to the synthesis of melanin (MEL). Oxidative bursts during the melanin-synthesis pathway can also create ROS that act as antimicrobials, which may also lead to priming, or antioxidant enzymes up-regulation (shown in bold letters and dark lines).

This integrative multivariate approach to tracking coral performance was utilized to test specific hypotheses regarding targeted response variables and the linkage between environmentally-driven physiological, symbiotic, and cellular (i.e., antioxidants, immunity) legacies as mechanisms underpinning bleaching outcomes (Palmer et al., 2010; Palmer & Traylor-Knowles, 2012; Venn, Loram, & Douglas, 2008). Both the host and symbiont have enzymatic defenses to mitigate oxidative stress (e.g., peroxidase, catalase, superoxide dismutase) (Venn et al., 2008) caused by host (e.g., heat-damaged mitochondrial membranes, Dunn et al., 2012) and symbiont (e.g., damage to the D1 protein of photosystem II (PSII)) (Jones, Hoegh-Guldberg, Larkum, & Schreiber, 1998; Lesser, 1997) sources, ultimately leading to apoptosis and dysbiosis when severe enough (Weis, 2008) (Figure 1b). Thermal stress also triggers host immune responses (Mydlarz, Couch, Weil, Smith, & Harvell, 2009; Mydlarz et al., 2010; Pinzón et al., 2015), and higher constitutive immunity (i.e., immune activity necessary for cellular homeostasis) reduces coral thermal sensitivity and disease susceptibility (Palmer et al., 2010). In addition, the melanin-synthesis pathway is an important component of host immunity that is active in wound healing and pathogen invasion while also serving as a photoprotectant that may mitigate bleaching stress (Mydlarz & Palmer, 2011; Palmer et al., 2010) (Figure 1b). However, corals may mount different cellular mechanisms to combat bleaching due to energetic requirements (Fuess, Mann, Jinks, Brinkhuis, & Mydlarz, 2018; Palmer, 2018b; Pinzón et al., 2015), stress frequency (i.e, acute, chronic, repeated stress) (Ainsworth et al., 2016; Schoepf et al., 2015), or as a function of histories of biotic and abiotic challenges that modulate constitutive immunity (Mydlarz et al., 2009; Palmer, 2018b; Wall et al., 2018). For instance, there is evidence corals may rely on different antioxidant and immunity mechanisms based on their environmental histories, as lower melanin synthesis and higher antioxidant activity was observed in heat-stressed corals from a reef with high pCO_2_-variability compared to low pCO_2_-variability (Wall et al., 2018). Thus, pairing the measurement of bleaching metrics like cell density and chlorophyll concentrations with immune and antioxidants provides a powerful and tractable approach within a multivariate integrative framework.

The Hawaiian Islands, once thought to be a bleaching refuge for corals (Jokiel & Brown, 2004), experienced severe bleaching in 2014 and 2015 across the populated Main and remote Northwestern Hawaiian Islands (Bahr, Rodgers, & Jokiel, 2017; Couch et al., 2017). In Kāne‘ohe Bay, O‘ahu, Hawai‘i, 7.1 degree heating weeks (DHW) were observed in 2014 and 10.2 DHW in 2015, beginning in August and July, respectively (*see* Methods), resulting in 45 ± 2% (mean ± SE) bleaching in the first event and 30 ± 4% bleaching in the second (Bahr et al., 2017). An ecologically dominant reef coral, *Montipora capitata* (Dana 1846), showed significant bleaching in both events (Figure 2a), with differences in bleaching responses observed at the site and event level.

**Figure 2.**
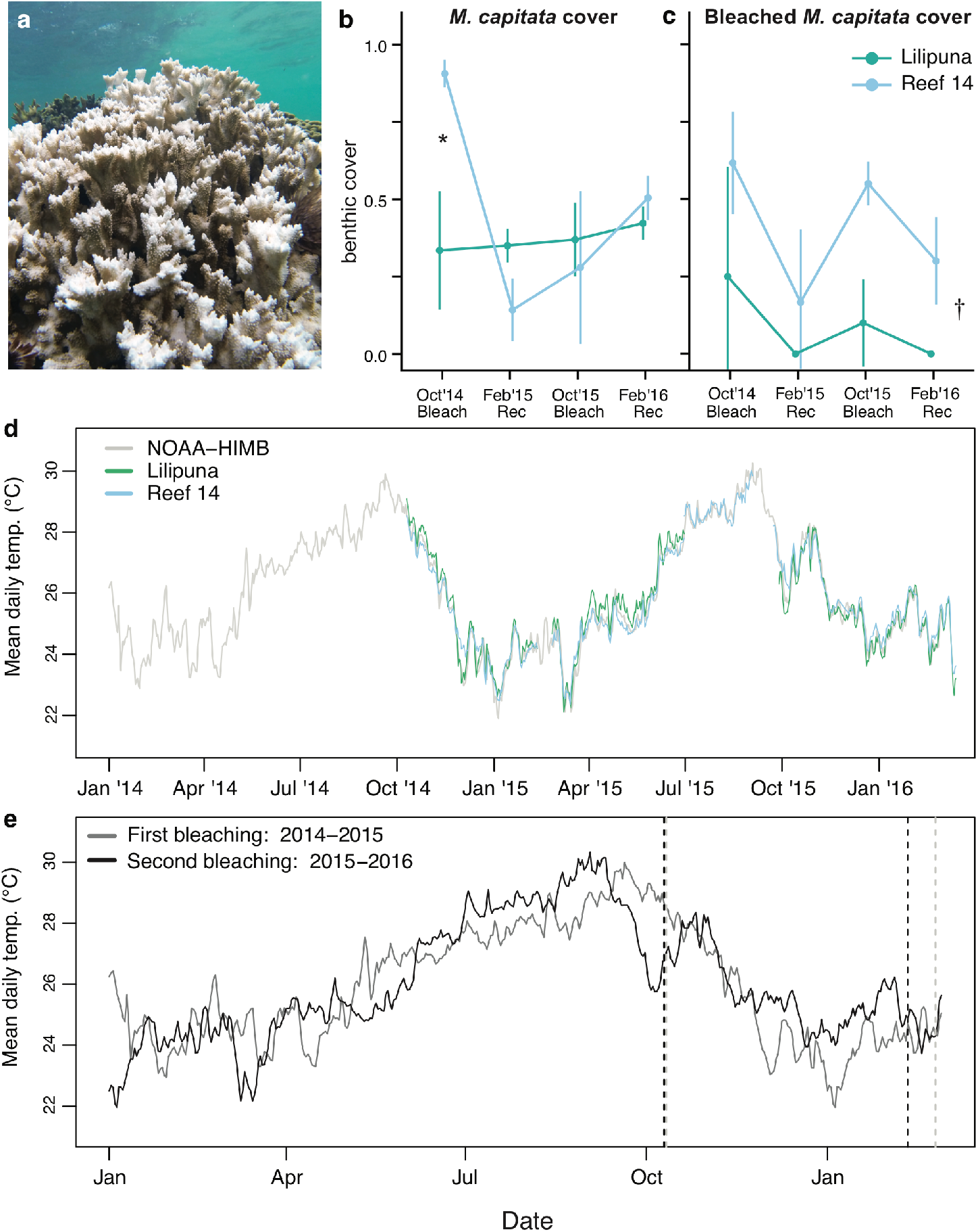
*Montipora capitata* cover and bleaching patterns at two reefs in relation to thermal regimes. (a) A bleached *M. capitata* colony and (b) benthic surveys of absolute *M. capitata* at two sites (Lilipuna and Reef 14) during and after thermal stress in 2014-2015 and 2015-2016. (c) The proportion of bleached *M. capitata* colonies scored during (October) and after (February) thermal stress (values are mean ± SD, *n* = duplicate transects). Symbols indicate differences among sites within a period (*) and among sites (†). (d) Overlay of mean daily temperatures at Lilipuna and Reef 14 and the NOAA-HIMB Moku o Lo‘e buoy from January 2014 - January 2016 and (e) a comparison of temperature ramping and cooling between the first and second bleaching events relative to our collection dates. Vertical dashed lines indicate coral collections during bleaching and recovery periods in the first (*black*, 2014-2015) and second (*gray*, 2015-2016) events.

Using the opportunity presented by these natural bleaching events (Figure 2), we tracked coral percent bleaching and physiotypes using a multivariate trait-space approach. We quantified the response of *M. capitata* in Kāne‘ohe Bay to the back-to-back bleaching events of 2014 and 2015 (i.e., bleaching in October 2014, recovery in February 2015, bleaching in October 2015, and recovery in February 2016) at two Kāne‘ohe Bay reefs (Lilipuna and Reef 14) with contrasting environmental histories relating to seawater residence (10 – 20 d *vs*. >30 d), pCO_2_ variability (ca. 300 *vs*. 600 *μ*atm pCO_2_ diel flux), and proximity to shore (Drupp, De Carlo, Mackenzie, Bienfang, & Sabine, 2011; Drupp et al., 2013; Lowe, Falter, Monismith, & Atkinson, 2009).

We predicted that: (H1) immunity and antioxidant contributions to bleaching responses will influenced by site environmental histories and symbiont communities such that greater bleaching prevalence and attenuated immune responses will be observed in corals from the high variable-pCO_2_ site (Reef 14) and those association with thermally sensitive Symbiodiniaceae. We also expected (H2) immunity and antioxidant activity would not occur with the same magnitude or trajectory after repeated bleaching events due to legacy effects shaping acclimatory or stress response pathways over time. Considering the wide-ranging functions of melanin in host immunity (Palmer et al., 2008, 2010), we expected melanin synthesis to act as an acute and broad-spectrum defense in physiologically stressed corals, with antioxidants having a more specialized response after chronic or repeat bleaching stress. Finally, while the thermal tolerance of symbiont communities is vital to coral bleaching sensitivity, evidence for differential immunity or antioxidant capacities in coral holobionts associated with thermally tolerant symbiont species is lacking. Given that the coral *M. capitata* is known to be dominated by two genetically and physiologically contrasting Symbiodiniaceae, *Cladocopium* sp. and *Durusdinium glynnii* (Cunning et al., 2016; Wall, Kaluhiokalani, Popp, Donahue, & Gates, 2020), we predicted (H3) that holobionts harboring heat-tolerant *D. glynnii* would show higher immune and antioxidant responses linked to greater bleaching resistance. To test these hypotheses, we measured bleaching severity and recovery at each location using benthic surveys and measured coral traits in coral fragments, including: physiological measurements of cellular bleaching magnitude (symbiont density, areal- and cell-specific chlorophyll *a*, holobiont total protein and total biomass), and completed enzymatic assays for mechanisms contributing to bleaching resistance through coral host antioxidant capacity (peroxidase, catalase, superoxide dismutase), and the host immune response of the melanin cascade (prophenoloxidase, melanin). This approach allows for a more holistic description of coral physiotype that we tracked through two bleaching and recovery periods.

## 2. Materials and Methods

### 2.1 Site description

Coral bleaching and recovery at two reef systems: a fringing reef (hereafter, ‘Lilipuna’ [21°25′36.8′′N, 157°47′24.0′′W]) in southern Kāne‘ohe Bay adjacent to the Hawai‘i Institute of Marine Biology (HIMB) on Moku o Lo‘e, and an inshore patch reef (hereafter, ‘Reef 14’ [21°27′08.6′′N, 157°48′04.7′′W]) in central Kāne‘ohe Bay. These reefs sites were chosen due to their unique environmental history of seawater pCO_2_ and hydrodynamics (Drupp et al., 2013; Wall et al., 2018). Seawater pCO_2_ adjacent to the both locations (fringing reef at Lilipuna and the patch reef Reef 14) is comparable (ca. 450 *μ*atm), however, diel pCO_2_ flux is significantly higher (196 – 976 *μ*atm pCO_2_) in central Kāne‘ohe Bay near Reef 14 relative to the Bay’s southern basin proximate to Lilipuna (225 – 671 *μ*atm) (P. Drupp et al., 2011; Drupp et al., 2013).

### 2.2 Environmental monitoring

Temperature (Hobo pendants, ± 0.53 °C accuracy, 0.14 °C resolution, Onset Computer Corp., Bourne, MA) and photosynthetic active radiation (PAR) loggers (Odyssey, Dataflow Systems Limited, Christchurch, New Zealand) were placed at Lilipuna and Reef 14 at a depth of 1 m. Loggers were periodically removed from the reef, cleaned, and replaced monthly. Temperature and PAR were recorded continuously at 15 min intervals from October 2014 – February 2016, however, gaps in environmental data do exist as a consequence of instrument failure and loss. Gaps in logger temperature data were supplemented with NOAA temperature data from the Moku o Lo’e station at HIMB (NOAA, 2019). PAR loggers were calibrated against a LI-1400 quantum meter (Li-Cor, Lincoln, Nebraska, USA) attached to a cosine LI-192 underwater quantum sensor. Temperature loggers were calibrated with a certified digital thermometer (5-077-8, ± 0.05 °C accuracy, Control Company, USA) and cross-calibrated against each other for standardization. Degree heating weeks (DHW) for the southern portion of Kāne‘ohe Bay where our corals were collected were calculated in R using *in situ* Moku o Lo‘e temperature data (NOAA, 2019), with the difference between mean half-week temperatures (i.e., mean hourly temp over 3.5 d) and the maximum monthly mean temperature of 27.7 °C, (Jokiel & Brown, 2004)] to determine ‘hotspots’. DHW were determined as the number of hotspots >1 across a rolling 12 weeks window (i.e., 24 half-weeks) (NOAA, 2020). DHW for windward O‘ahu and the Hawaiian Islands during the 2014 and 2015 bleaching events have been previously reported (Bahr et al., 2017; Sale, Marko, Oliver, & Hunter, 2019). The purpose of our calculations are to quantify the incurred heat stress using empirical calculation of DHW from temperature data collected proximate to our two reef locations.

### 2.3 Benthic surveys and coral collections

Four sampling periods were identified as corresponding to a ‘bleaching period’ following the point of maximum thermal stress (10 October 2014 and 12 October 2015) and a post-bleaching ‘recovery period’ approximately 4 months after peak seawater warming (11 February 2015 and 26 February 2016). In each time period, benthic surveys were conducted at each reef site using a 20 m transect and a line-point-intersect at 1 m intervals. Transects were positioned parallel to natural contours of the reef, being the north-south axis of the fringing reef (Lilipuna) and the east-west axis of patch reef (Reef 14). At each reef, transects (*n* = 2) were placed along the reef crest (1 – 2 m), and benthic community cover was recorded at the species level for reef corals (*Montipora capitata, Pocillopora* spp. [*P. acuta*, *P. damicornis*]*, Porites compressa*), and either crustose coralline algae (CCA), macroalgae, or sand/bare/turf. For corals, bleaching state was quantified categorically, being either non-bleached (i.e., appearing fully pigmented), or bleached (i.e., exhibiting degrees of tissue paling/pigment variegation or being wholly white). *M. capitata* cover was calculated as the proportion of total benthic cover, and *M. capitata* bleaching extent was calculated as proportion of total *M. capitata* colonies bleached. The relative position of transects at each reef was recorded to allow for repeat surveys within the same general location at each reef across survey periods.

In each sampling period, coral branch tips (<4 cm length) were collected from forty *M. capitata* coral colonies (*n* = 1 fragment colony^−1^) along the reef crest at each reef site at a depth of ca. 1 m (State of Hawai‘i Department of Land and Natural Resources, Special Activity Permit 2015-17 and 2016-69). Immediately post collection, corals were snap-frozen in liquid nitrogen and returned to HIMB and stored at −80 °C. While remaining frozen, each colony was split in half along its longitudinal axis. One-half of each coral fragment was stored at HIMB (−80 °C) until processing for physiological assays and qPCR. The corresponding fragment-halves were used for immunological assays and were shipped to the University of Texas at Arlington using a dry-shipper charged with liquid nitrogen.

### 2.4 DNA extraction and symbiont community analysis

Symbiodiniaceae DNA was extracted by adding an isolate of coral tissue (500 *μ*l) to 500 ul DNA buffer (0.4 M NaCl, 0.05 M EDTA) with 2% (w/v) sodium dodecyl sulfate, following a modified CTAB-chloroform protocol (Cunning et al., 2016) (dx.doi.org/10.17504/protocols.io.dyq7vv).

To determine the dominant genera of Symbiodiniaceae hosted by each *M. capitata* fragment, a quantitative PCR (qPCR) (Cunning et al., 2016) was used to amplify Symbiodiniaceae actin gene loci, with the corresponding number of actin gene copies in the Symbiodiniaceae genera *Cladocopium* and *Durusdinium* (namely, ITS2 types C31, with C17 and C21 and D1-4-6 [*Durusdinium glynnii* (Wham, Ning, & LaJeunesse, 2017)]) known to be numerically dominant in Kāne‘ohe Bay *M. capitata* (Cunning et al., 2016). Specificity of genus-level primers have been previously validated using a combination of Symbiodiniaceae internal transcribed spacer (ITS2) region of rDNA and actin gene sequencing (Cunning & Baker, 2013). Duplicate qPCR reactions (10 *μ*l) were run for each coral sample using a StepOnePlus platform (Applied Biosystems) set to 40 cycles, a relative fluorescence (ΔR_n_) threshold of 0.01, and internal cycle baseline of 3 – 15. Symbiont genera detected in only one technical replicate were considered absent. The relative abundance *of Cladocopium and Durusdinium* symbionts (i.e., C:D ratio) in each sample was determined from the ratio of amplification threshold cycles (C_T_) for *Cladocopium* and *Durusdinium* (i.e., C_T_^C^, C_T_^D^) using the formula C:D = 2^(C_T_^C^ - C_T_^D^), where genus-specific C_T_ values are normalized according to gene locus copy number and fluorescence intensity (Cunning et al., 2016). Coral colonies were determined to be *Cladocopium*- or *Durusdinium*-dominated based on numerical abundance (>0.5 proportion) of each genus from qPCR analysis (Innis, Cunning, Ritson-Williams, Wall, & Gates, 2018).

### 2.5 Physiological metrics

The extraction and processing of coral and symbiont tissues was performed following established methods (summarized in Wall et al., 2018). Briefly, coral tissue was removed from the skeleton using an airbrush filled with 0.2 *μ*m filtered seawater, yielding ~10 – 30 ml of tissue slurry.

Extracted tissues were briefly homogenized and subsampled for the following physiological metrics: symbiont cell densities, chlorophyll *a* concentrations, protein biomass, and the total organic biomass determined from the ash-free dry weight (AFDW) of coral + algae tissues. Coral tissue slurries were stored at −20 °C.

All physiological metrics were normalized to the surface area (cm^2^) of coral skeleton using the paraffin wax-dipping technique (Stimson & Kinzie, 1991). Symbiont cell counts were measured by replicate cell counts (*n =* 6 – 10) on a haemocytometer, and expressed as symbiont cells cm^−2^. Chlorophyll *a* was quantified by concentrating algal cells through centrifugation (3,000 × *g* for 3 min) and extracting pigments in the algal pellet in 100% acetone for 36 h in darkness at −20 °C. Spectrometric absorbance were measured (*λ* = 630 and 663 nm) using a 96-well quartz plate with two technical replicates; chlorophyll concentrations quantified using the equations for dinoflagellates (Jeffrey & Humphrey, 1975), normalized for path length and expressed as ug chlorophyll *a* cm^−2^ and pg chlorophyll *a* symbiont cell^−1^. Total protein concentration (soluble + insoluble in the holobiont) was quantified using the Pierce BCA (bicinchoninic acid) Protein Assay Kit (Pierce Biotechnology, Waltham, MA). Protein solubilization was achieved by adding 1 M NaOH and heating (90 °C) for 60 min, followed by the neutralizing to ca. pH 7.5 with 1 N HCl. Protein was measured spectrophotometrically (*λ* = 562 nm) in a 96-well plate with three technical replicates against a bovine serum albumin standard and expressed as mg protein cm^−2^. Total fraction of organic biomass was measured by drying a subsample of coral tissue at 60 °C (48 h) in pre-burned aluminum pans followed by burning in a muffle furnace (450 °C) for 4 h. The difference between the dried and burned masses is the AFDW and expressed as mg cm^−2^.

### 2.6 Immunity and oxidative stress metrics

Immunology and oxidative stress metrics were determined using previously published protocols for coral host tissues (Mydlarz et al., 2009; Mydlarz & Palmer, 2011; Palmer et al., 2011; Wall et al., 2018). A 3 – 4 ml aliquot of coral tissue slurry was obtained by airbrushing with our coral extraction buffer at pH 7.0 (100 mM TRIS buffer + 0.05 mM dithiothreitol). Tissue was homogenized for 1 min on ice (hand held homogenizer, Powergen 125, Fisher Scientific, Waltham, MA), and 1 ml of the resulting slurry was freeze-dried for 24 h (VirTis BTK freeze-dryer, SP Scientific, Warminster, PA) and used for melanin concentration estimates. The remaining slurry was centrifuged at 4 °C at 2,500 × *g* (Eppindorf 5810 R centrifuge, Hamburg, Germany) for 5 min to remove cellular debris and most Symbiodiniaceae cells to achieve a host-enriched cell-free extract. All colorimetric measurements were calculated using a Synergy 2 multi-Detection microplate reader with Gen5 software (BioTek, Winooski, VT, USA). All assays were run in duplicate or triplicate on separate 96-well microtiter plates. Total protein concentration of each coral cell-free extract was determined using the RED660 protein assay (G Biosciences, Saint Louis, MO) with a bovine serum albumin standard curve.

To determine the concentration of melanin within each sample, the freeze-dried aliquots of weighed and dried tissue were gently vortexed with 400 *μ*l of 10 M sodium hydroxide (NaOH) and left to extract for 48 h. Samples were bead-beaten with 1 mm glass beads and then vortexed for 10 s and then centrifuged at 7,000 × *g* for 5min. For each sample, 65 *μ*l of supernatant were aliquoted in duplicate into a 96-well g-area microtiter plate (Greiner Bioone, Monroe, NC, USA). The plates were read at an absorbance of 490 nm and the concentration of melanin was determined using a standard curve (0 – 2 mg) of commercial melanin (Sigma-Aldrich, St. Louis, MO) dissolved in 10 M NaOH for 48 h and treated the same as the samples on each microtiter plate. Data presented are converted to mg melanin normalized to mg of tissue (Fuess et al., 2018).

PPO activity was determined for each sample using duplicate 20 *μ*l aliquots of coral cell-free extract in clear 96-well microtiter plates, with 50 *μ*l of 10 mM phosphate buffered saline at pH 7.0. To each well, 20 *μ*l of trypsin (0.2 mg ml^−1^ concentration in deionized filtered water) was added to activate PPO and the reaction was initiated by the addition of 20 *μ*l of 25 mM L-1,3-dihydroxyphenylalanine (L-DOPA; Sigma-Aldrich). The absorbance at 490 nm was measured over 25 min and the change in absorbance during the linear portion of the reaction (typically 10 – 15 min) was normalized to mg protein and time for each sample (ΔAbs490 nm mg protein^−1^ min^−1^) (Mydlarz & Palmer, 2011; Fuess et al., 2018).

Coral host oxidative stress was determined by measuring the scavenging activity of the coral cell-free extracts to different substrates comparable to the antioxidants: peroxidase (POX), catalase (CAT), and superoxide dismutase (SOD). Peroxidase activity (EC 1.11.1.7) was determined for each sample using 10 *μ*l aliquots of coral cell-free extract, in duplicate within 96-well microtiter plates, diluted with 50 *μ*l phosphate buffer (pH6.0). To each well, of 25 *μ*l of 25 mM guaiacol in 10 mM PBS (pH 6.0) was added and the reaction was initiated with the addition of 12.5 *μ*l of 25 mM H_2_O_2_ high purity 30% hydrogen peroxide (H_2_O_2_; Sigma Aldrich). Guaiacol is a phenolic compound with scavenging properties that is commonly used as a substrate for peroxidase activity (Mydlarz & Harvell, 2007). The absorbance was read at 470 nm every min for 15 min and the change over time was calculated for the linear part of the reaction (0 – 10 min) and normalized to mg protein in each sample the units for peroxidase activity are (ΔAbs470 nm mg protein^−1^ min^−1^) (Mydlarz & Harvell, 2008).

Catalase activity (EC 1.11.1.6) was determined using the H_2_O_2_ depletion assay using 10 *μ*l coral cell free extract and 50 *μ*l of 10 mM of pH 6.0 phosphate buffered saline (PBS) in duplicate wells of a UV-transparent 96-well microtiter plate. The assay was activated by the addition of 10 *μ*l 25 mM H_2_O_2_. The absorbance was measured at 240 nm over 15 min and the change in absorbance over time (e.g., initial - final) was calculated for the linear part of the reaction (typically 0 – 8 min) and converted to *μ*mol H_2_O_2_ using a standard curve of known concentrations, normalized per mg protein. CAT activity is presented as *μ*mol H_2_O_2_ scavenged mg protein^−1^ min^−1^ (Palmer et al., 2011).

Superoxide dismutase (EC 1.15.1.1) activity was calculated using the SOD Assay Determination Kit-WST (Fluka, Switzerland) which includes Dojindo’s highly water-soluble tetrazolium salt, WST-1(2-(4-iodophenyl)-3-(4-nitrophenyl)-5-(2,4-disulfophenyl)-2H-tetrazolium, monosodium salt), that produces a water-soluble formazan dye with an absorbance peak at 450 nm upon reduction with superoxide anion. 10 *μ*L of extract was incubated with WST-1 and xanthine oxidase, which produces superoxide anion. Inhibition of absorbance at 450 nm (scavenging of the superoxide anion) was monitored in wells containing coral extracts and SOD standards and compared to untreated samples. One SOD unit of activity (U) equates to 50% inhibition of superoxide anion, and data are presented as SOD U mg protein^−1^ (Krueger et al., 2015).

### 2.7 Statistical analysis

Permutation analysis of variance (PERMANOVA) and non-metric multidimensional scaling (NMDS) were performed using a balanced matrix of all physiology and immunity/antioxidant responses (*n =* 10 responses) in the package *vegan* (Oksanen et al., 2019). PERMANOVA used a scaled and centered matrix with Euclidean calculations of pairwise distances using *adonis2*. The NMDS data matrix was double standardized using a Wisconsin and square root transformation with a euclidean distance using *metaMDS*. Response variables showing significant correlations (*p*<0.05) with NMDS axes were plotted as vectors using envfit command in *vegan*. Coral ‘physiotypes’ were defined by convex hulls, with borders defined by the range of points in the multivariate trait space (i.e., NMDS1 and NMDS2). Physiotypes were grouped categorically by periods, or the interactions of Site, Symbiont, and Period. Physiotype centroids used in trajectory plots were calculated as the mean of NMDS1 and NMD2 for each category.

Ecological benthic data (*M. capitata* total cover and bleached cover) were tested in a linear model with Periods (two bleaching and two recovery events) and Sites (Lilipuna, Reef 14) as fixed effects. Physiology, antioxidant, and immunity response variables were analyzed using a linear model with Periods, Sites, and Symbiont community composition (*Cladocopium*- or *Durusdinium*-dominated) as fixed effects. Normal distribution and equal variance assumptions of ANOVA were examined by graphical representation of residuals and quantile:quantile plots. Where assumptions were not met, Box-Cox tests were performed (Box & Cox, 1964) and data transformations applied in the package *MASS* (Venable & Ripley, 2002). Analysis of variance tables for linear models were generated using type-II sum or squares with multiple comparisons using the package *emmeans* (Lenth, 2019). All analyses were performed in *R* version 3.6.1 (R Core Team, 2019). All data and code to generate figures and perform analyses are archived and openly available at Github (https://github.com/cbwall/Gates-Mydlarz-bleaching-recovery/releases/tag/v4).

## 3. Results

Coral cover at the two sites ranged from (mean ± SD) 63 ± 11% to 93 ± 11% from 2014-2016. Greater total bleached coral cover was observed in the first bleaching event (62 – 75 ± 9%) relative to the second (43 – 55 ± 16%) (i.e., during thermal stress) (Figure 2). Mean *M. capitata* cover was stable across time at Lilipuna (~40%) but was much more variable at Reef 14, falling from 91 ± 4% in October 2014 to 51 ± 7% in February 2016. In the two bleaching events (October 2014 and 2015) greater bleaching of *M. capitata* corals was observed at Reef 14, compared to Lilipuna (*p*=0.020).

PERMANOVA results testing effects of Period (bleaching event), Site (Lilipuna *vs*. Reef 14) and dominant Symbiodiniaceae species (*Cladocopium* sp. or *D. glynnii*) revealed significant differences of all main effects (*p*<0.001) and the interaction of Period-by-Site (*p*<0.001) and Period-by-Symbiont community (*p*<0.001) (Figure 3 and 4, Table 1). In each of the four periods, coral physiotypes occupied unique positions in multidimensional trait space (Figure 2), with the location and trajectories of coral physiotypes in each period principally being a function of symbiont community, and to a lesser extent, site of collection (i.e., environmental history and physiological legacy) (Figure 3, 4).

**Table 1.** Results of PERMANOVA testing the effects of repeated bleaching and recovery periods on *Montipora capitata* corals hosting two distinct symbiont communities at two reef locations.

| *Factor* | *df* | *SS* | *R*<sup>2</sup> | *F* | *p* |
| --- | --- | --- | --- | --- | --- |
| Period | 3 | 855.740 | 0.297 | 48.491 | **<0.001** |
| Site | 1 | 60.996 | 0.021 | 10.369 | **<0.001** |
| Symbiont | 1 | 202.349 | 0.070 | 34.399 | **<0.001** |
| Period × Site | 3 | 69.003 | 0.024 | 3.910 | **<0.001** |
| Period × Symbiont | 3 | 60.101 | 0.021 | 3.406 | **<0.001** |
| Site × Symbiont | 1 | 6.861 | 0.002 | 1.166 | 0.293 |
| Period × Site × Symbiont | 3 | 19.048 | 0.007 | 1.079 | 0.360 |
| Residual | 273 | 1605.902 | 0.558 |  |  |
| Total | 288 | 2880.000 | 1.000 |  |  |

*Period* = sequential bleaching and recovery events from October 2014 – February 2016, *Site* = Lilipuna or Reef 14, *Symbiont* = *Cladocopium* sp. or *Durisdinium glynnii* dominated symbiont community. *SS* = sum of squares; *df* = degrees of freedom; bold *p* values represent significant effects ( $p < 0.05$ ).

**Figure 3.**
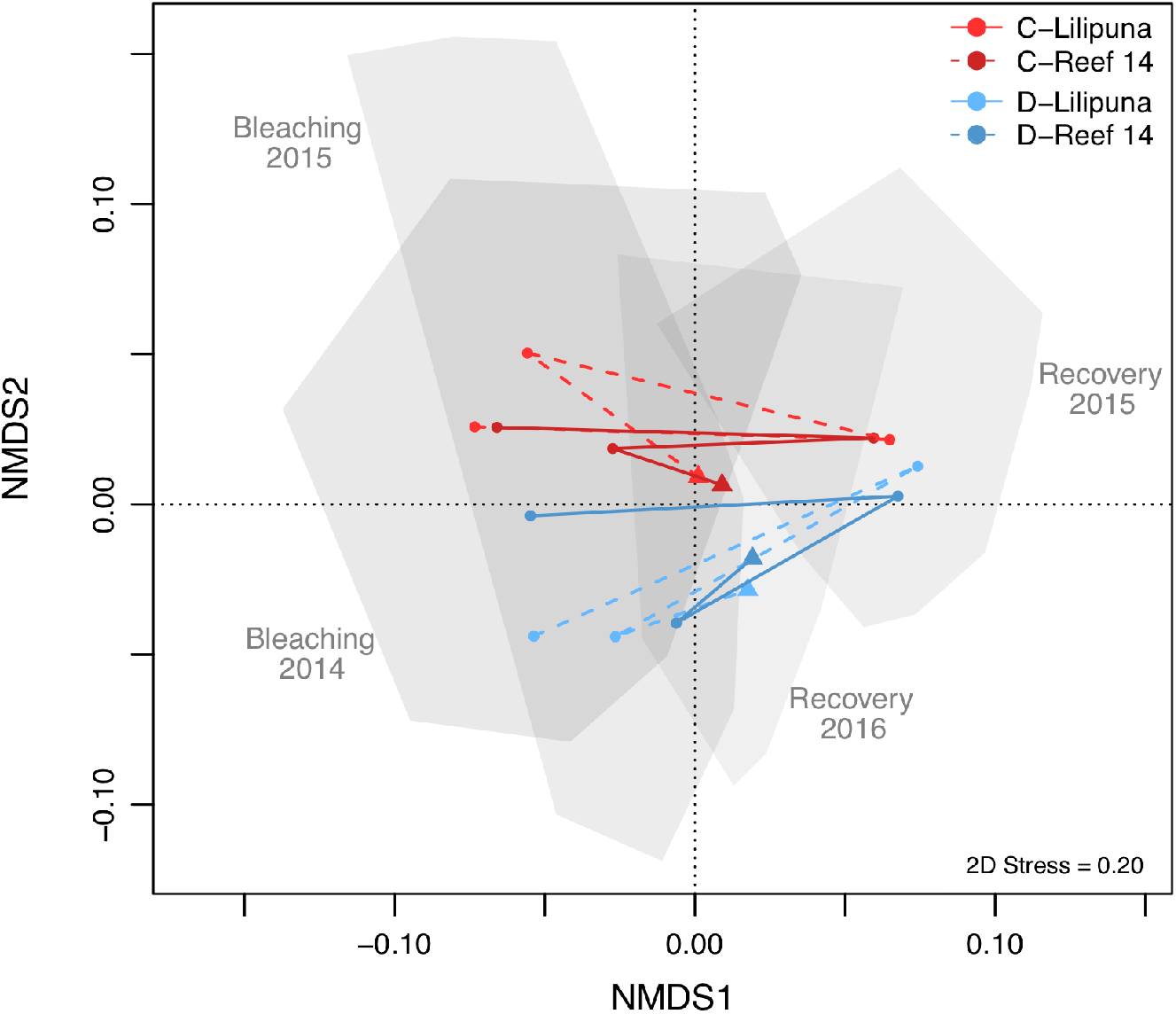
Non-metric multidimensional scaling (NMDS) analyses of coral performance envelopes during bleaching stress and post-bleaching recovery. *Montipora capitata* corals dominated by *Cladocopium* sp. (red) or *Durusdinium glynnii* (blue) symbionts from two sites (*solid lines* Lilipuna, *dashed lines* Reef 14). Convex hulls represent the coral physiotype (i.e., NMDS point clusters) of all corals in each time period, with points indicating mean centroids of physiotype polygon for each group. Lines show trajectories of mean centroids for each group across the four periods (Bleaching 2014, Recovery 2015, Bleaching 2015, Recovery 2016) and triangle arrowheads indicate trajectory termini.

**Figure 4.**
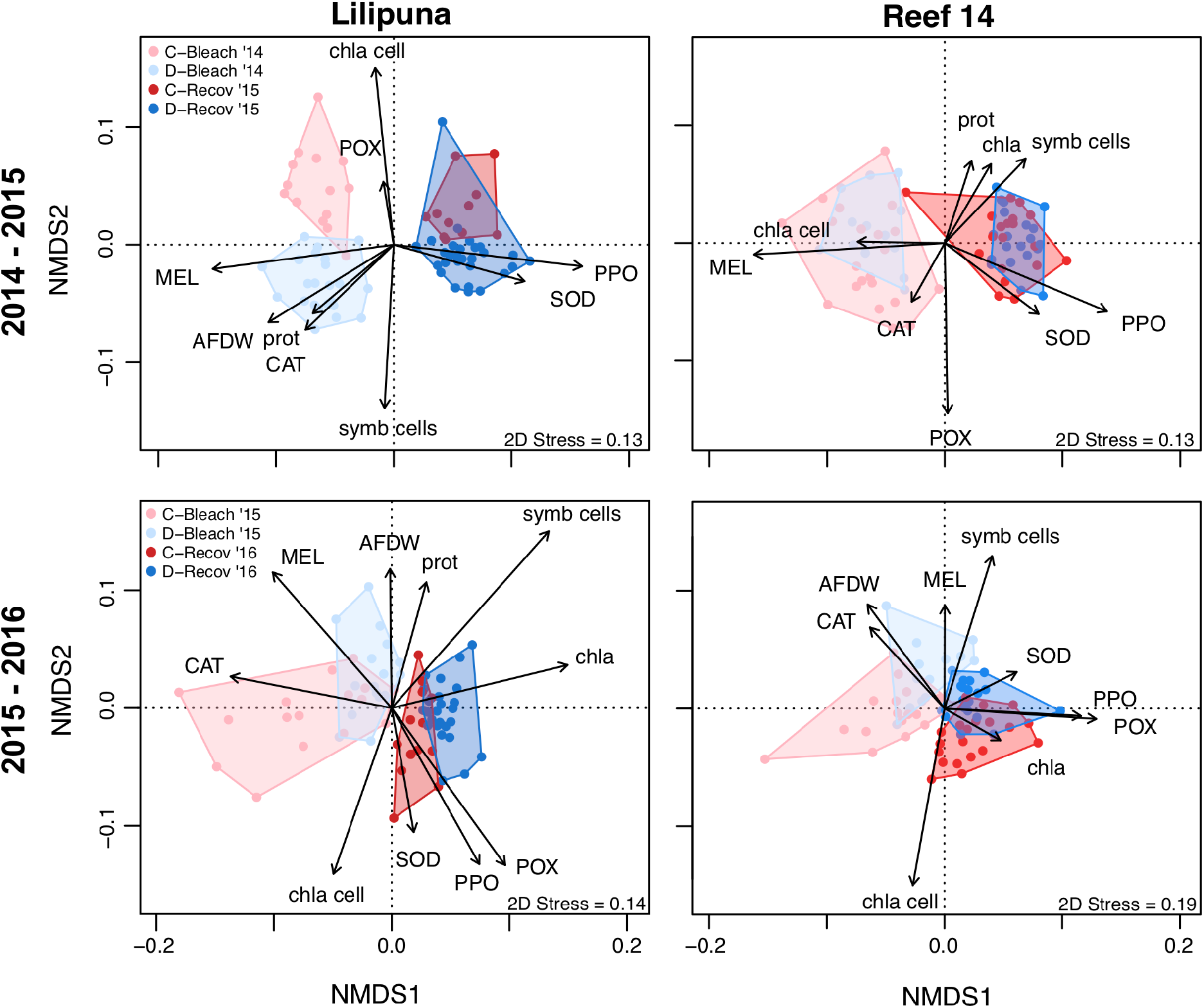
Non-metric multidimensional scaling (NMDS) analyses identify coral physiotypes separated by dominant symbiont type (colors), bleaching-recovery periods (lighter and darker shades), sites (columns), and events (rows). *Montipora capitata* corals dominated by *Cladocopium* sp. or *Durusdinium glynnii* symbionts from Lilipuna (left panel) and Reef 14 (right panel) during bleaching (Bleach) and recovery (Recov) periods. Biplot vectors (black arrows) represent significant physiology and immunity responses (*p* < 0.05) according to squared correlation coefficients (*r*^2^). AFDW = ash-free dry weight biomass (mg gdw^−1^), CAT = catalase, MEL = melanin, POX = peroxidase, PPO = prophenoloxidase, SOD = superoxide dismutase, chla = chlorophyll *a μg* cm^−2^, chla cell = chlorophyll *a* per symbiont cell, prot = protein mg cm^−2^, symb cells = symbionts cm^−2^.

Physiotypes of bleached corals relative to recovered corals showed stronger separation in 2014-2015, which was apparent at both reef sites (Figure 3, top row) and paralleled patterns of greater bleaching prevalence in the first-bleaching event (Figure 2c). Similarly, site effects on coral physiotypes were most pronounced in the first bleaching event (Table 1, Figure 4). Despite greater bleaching at Reef 14 in October 2014 (Figure 2b), shifts in bleaching and recovery performance envelopes were most distinct at Lilipuna (Figure 4). At the physiological level, separation of physiotypes during and after thermal stress was driven by bleaching sensitivity and symbiont cells/chlorophyll concentrations (*p*<0.001, Figure 4 & S2, Table S1). “Bleached” physiotypes observed in periods of temperature stress (October 2014 and 2015) generally aligned with colonies hosting the more thermally sensitive *Cladocopium* sp., although in the first event coral physiotypes at Reef 14 symbiont communities did not separate according to symbiont communities (Figure 4). Nevertheless, across all sampling periods *M. capitata* colonies dominated by *Durusdinium* symbionts had higher symbiont cell densities, less variable areal chlorophyll concentrations, and lower chlorophyll per symbiont cell (i.e., pg chlorophyll cell^−1^) compared to colonies dominated by *Cladocopium* symbionts (*p*<0.001, Figure S2). While *Cladocopium* chlorophyll cell^−1^ oscillated across bleaching-and recovery periods, *Durusdinium* pg chlorophyll cell^−1^ remained stable (*p*=0.010) and was also slightly higher at Reef 14 relative to Lilipuna across all timepoints (*p*=0.001, Table S1). Coral protein and total biomass was variable across the study, but each showed positive correlations with *Durusdinium*-dominated physiotypes from Lilipuna in bleaching and recovery periods (Figure 4). Overall, protein was 9% higher in *Durusdinum-*dominated colonies (*p=*0.031) and 32% higher in Lilipuna corals during first bleaching (October 2014) but equivalent at all sampling points thereafter (*p*=0.002). Total biomass was lower during recovery periods (*p*<0.001) in addition to being 20 – 40% higher at Lilipuna compared to Reef 14 in all periods (except in February 2015 recovery) (*p*<0.001) and ~30% higher in *Durusdinium*-dominated colonies during the second bleaching, but equivalent across all colonies in other periods (*p*=0.012) (Figure S2).

Immunity and antioxidant metrics (i.e., MEL, PPO, POX, CAT, SOD) differed through time in response to repeat bleaching and recovery (Figure 3, Figure 4) (*p*<0.001), while the environmental influence from the Sit and Symbiont communities were less pronounced (Figure S2, S3, and Table S1, S2). In the first event, corals responded to thermal stress by increasing melanin synthesis (*p*<0.001) (with corresponding declines in PPO precursors) and increasing catalase (*p*<0.001). During recovery, prophenoloxidase (*post-hoc*: *p*<0.001) and superoxide dismutase increased (*post-hoc*: *p*<0.001) as melanin and catalase declined (Figure 4, 5, S2, & S3, Table S1, S2). Notably, the melanin pathway increased in all corals in the 2014-bleaching event regardless of bleaching sensitivity (i.e., loss or retention of symbionts/chlorophyll), or symbiont community, and was a significant cellular response that shaped coral physiotypes in the first bleaching event (Figure 5, first row). Corals in the second bleaching experienced a 6-fold increase in melanin (October 2015 *vs*. February 2015) along with corresponding reductions in prophenoloxidase, however, melanin activity was significantly less compared to the high melanin activity observed in the first bleaching period (0.015 *vs*. 0.003 mg melanin mg tissue^−1^ in first-versus second-bleaching period). Conversely to lower melanin in the second bleaching was a peak in catalase activity, which reached its highest level in 2015 bleaching (*post-hoc*, *p*<0.001) and was most pronounced in corals from Lilipuna compared to Reef 14 (*post-hoc*, *p*<0.001) (Table S2). The peak in catalase during 2015 bleaching corresponded with the lowest observed peroxidase activity, particularly for *Cladocopium*-dominated colonies (*post-hoc*, *p*=0.007). Subsequently in the 2016 recovery period, catalase declined by ~70% and peroxidase activity doubled, reaching peak activity similar to those observed in time periods. In contrast to other immunity and antioxidant responses, superoxide dismutase showed a unique trajectory, increasing progressively through time with each sampling period, being at its highest constitutive activity after repeat bleaching in the 2016 recovery, which was double that observed in the first bleaching period (October 2014) (*p*<0.001). Similar to catalase, superoxide dismutase was also slightly higher in corals from Lilipuna compared to those at Reef 14 (*p*=0.028).

**Figure 5.**
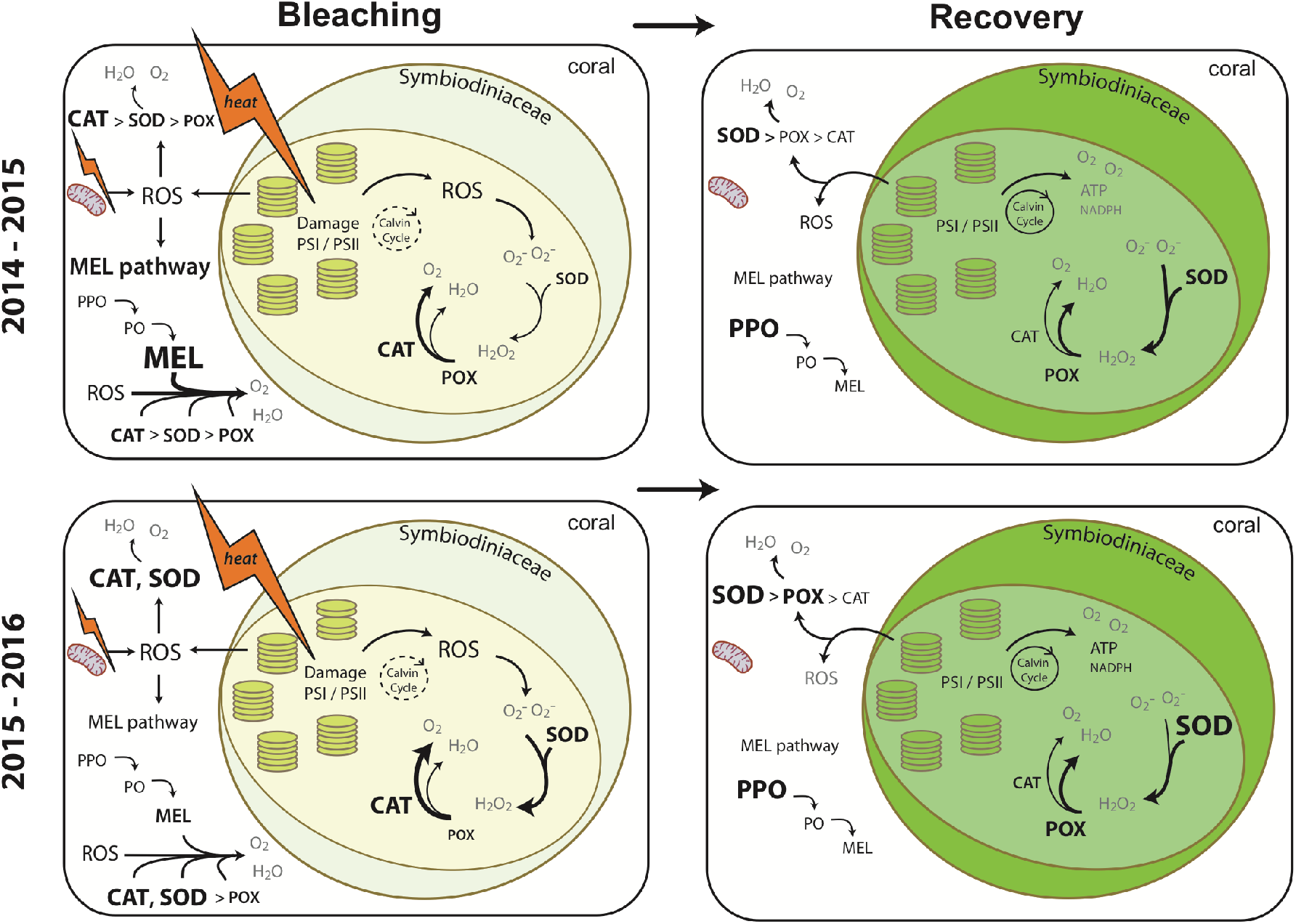
Schematic of mechanistic coral responses to repeat bleaching and recovery leading to shifted physiotypes. In the first event (*top row*) thermal stress leads to a substantial increase in MEL along with modest spikes in CAT and SOD. Corals in the first recovery showed increases in PPO (as the precursor to the melanin-pathway) as well as SOD and POX, while CAT activity declined. In the second event (*bottom row*), CAT and SOD spike with modest contributions of MEL and a general decline in POX. Corals in the second recovery had the highest levels of SOD across all time points, an increase in POX and a sharp decline in CAT (*see* Figure 4 for abbreviations).

## 4. Discussion

Ocean thermal anomalies are reshaping the structure and function of coral reefs the world over. Three mass bleaching events occurred since 2014 (Hughes et al., 2017), and climate models predict that by mid-century bleaching will be an annual phenomena (van Hooidonk, Maynard, & Planes, 2013). The stress responses of corals to ocean warming is based on a network of dynamic interactions at biological and environmental levels, that can influence how they respond to the physiological challenges posed by a warming planet (Suggett & Smith, 2019). Accordingly, there is a need for holistic measures of coral response that incorporate the influence of and environmental history (Palumbi et al., 2014) on physiological legacy effects. For example assessment of symbiont community (Suggett, Warner, & Leggat, 2017) and cellular memory (Brown, Dunne, Edwards, Sweet, & Phongsuwan, 2015) in the coral holobiont during and after thermal stress, to test for shifting physiological status that may ameliorate or exacerbate coral bleaching. Thus, physiotype tracking – or multivariate physiological and molecular time series data – provide the capacity for identification of the cellular mechanisms that contribute to resilience, acclimatization, and/or resistance that can allow for better predictions of coral ecological responses to intensifying climate change and other regional disturbances (Vercelloni et al., 2020).

Symbiosis – via differential performance of genera and types (Sampayo et al., 2008; Stat et al., 2008) and the potential for switching and shuffling (Buddemeier & Fautin 1993; Baker et al., 2003) – provide one clear example of how environmental history can shift performance baselines. In this study, the role of dominant Symbiodiniaceae on holobiont physiotype was present, but the symbiont community effects were often Period- or Site-dependent. The high symbiont and chlorophyll *a* densities in *M. capitata* colonies dominated by *D. glynnii* across Periods aligns with the paradigm of high thermotolerance within the genus *Durusdinium* (Cunning et al., 2016; Lesser, Stat, & Gates, 2013; Silverstein, Cunning, & Baker, 2017; Wham et al., 2017). However, antioxidant and immunity metrics were equivalent in colonies differing in dominant symbiont partner, demonstrating negligible effects on antioxidant and immune traits counter to the expectation that these would be modulated by symbiont-derived bleaching resistance. In fact, this similarity was despite a greater loss of symbiont cells and photopigmentation in colonies dominated by *Cladocopium* relative to *Durusdinium*. Symbiont PSII photodamge is thought to lead the cascade of cellular events that culminate in bleaching (Weis, 2008). However, some Symbiodiniaceae (*Symbiodinium* sp., *Durusdinium trenchii*) have been shown to resist expulsion from the host despite incurring photodamage (Kemp et al., 2011; Silverstein et al., 2017). Therefore, host mechanisms regulating redox status may be equally important in coral thermal stress responses and decoupled from symbiont photophysiological function (Krueger et al., 2015). In our study, the similarities observed between the antioxidant and immune responses of *Cladocopium*- and *Durusdinium-*dominated *M. capitata* colonies during both bleaching and recovery periods implicate an integral role of host mechanisms in coral responses to bleaching and recovery despite the presence of functionally distinct symbiont communities.

Shifts in physiotypes across bleaching and recovery periods were influenced by symbiont communities and the physical environmental at each site, however, the significance of these predictors were not uniform across time and varied during first- and second-bleaching events. For instance, the effects of reef site were clear in the first event, possibly relating to greater bleaching in 2014, which reduced symbiont cells in both corals dominated by *Cladocopium* sp. and thermotolerant *D. glynii* at Reef 14. In the context of previous works, our data agrees with findings that DHW in Kāne‘ohe Bay were lower in the first bleaching event relative to the second (5 – 7 DHW [2014] *vs*. 10 – 12 DHW [2015]) (Bahr et al., 2017), while the percentage of bleached coral cover was higher in 2014 at our study sites (70% [2014] *vs*. 50% [2015]) and in Kāne‘ohe Bay as a whole (45 – 77% [2014] *vs*. 30 – 55% [2015]) (Bahr et al., 2017; Ritson-Williams & Gates, 2020) possibly due to greater cumulative temperature stress in 2014 (Bahr et al., 2017; Ritson-Williams & Gates, 2020). While this study and others agree that lower bleaching prevalence was observed in Kāne‘ohe Bay during the second bleaching event, we show this corresponded with attenuated Site × Symbiont effects on coral physiotypes (Figure 4 *bottom row*), suggesting a positive feedback between this interaction and bleaching severity. Therefore, the symbiont community harbored by corals is integral to bleaching responses and coral physiotypes (Suggett et al., 2017), but the relative importance of symbiont effects are also tempered by present environmental conditions and site-specific environmental histories.

Environmental conditions between Lilipuna and Reef 14 did not differ in terms of light availability, seawater temperature, or DHW (Figures 2 & S1). Nevertheless, the hydrodynamics between these reefs are substantial, producing considerable differences in seawater residence and pCO_2_ variability (Drupp et al. 2011, 2013). In addition, the fringing reef habitat of Lilipuna with a close proximity to shore and silt-dominated backreef benthos is in stark contrast to Reef 14, which is patch reef pinnacle in the middle of the Kāne‘ohe Bay lagoon. Together, the persistence of long-term hydrodynamic and biogeochemistry conditions between these reefs have manifested as physiological legacies in resident *M. capitata*, which we demonstrate influence the response of corals to regional bleaching events. In the laboratory, we showed *M. capitata* from Reef 14 exhibited greater antioxidant activity and *F_v_/F_m_* but lower melanin compared to Lilipuna colones (Wall et al., 2018). We hypothesize these patterns observed in laboratory and field studies may be underpinned by energetic limitations due to pCO_2_ histories. Immunity is energetically expensive (Palmer et al. 2018b) and studies show corals exposed to high-pCO_2_ upregulate genes for energy reserve metabolism (Vidal-Dupiol et al., 2013) and can have lower lipid biomass compared to corals at ambient-pCO_2_ (Wall, Mason, Ellis, Cunning, & Gates, 2017). In this case flow and pCO_2_-history may be an important factor in the physiological legacies of Kāne‘ohe Bay *M. capitata*, which had a greater impact on during the first bleaching event relative to the second.

Coral physiotypes are influenced by the physiological state and integrity of the coral-Symbiodiniaceae symbiosis, which includes (but is not limited to) pigmentation and symbiont densities. As expected, we observed symbiont loss/repopulation to drive changes in coral physiotypes between bleaching and recovery periods, and in support of our hypothesis, prior events had cumulative impacts on physiotype trajectories. Notably, coral physiotypes occupied distinct spaces in each sampling period and showed changing trajectory through time (2014-2016) with the terminus of the trajectory in 2016-recovery resulting in a coral trait space intermediate between prior bleaching and recovery periods (Figure 1). These trajectories may be driven by those responses active in either the first or second bleaching-recovery periods (such as melanin), or by changes in constitutive immune activity that are cyclical (e.g., PPO and CAT), or continuously increasing through time (e.g., SOD). For instance, changes in total biomass generally declined in the aftermath of bleaching, being lower in recovery periods, but did not follow patterns in symbiont or photopigment concentrations. Coral tissue biomass also increased through time and was highest in the second year (on average pooled across all samples). The progressive increases in constitutive antioxidant activity exemplified by superoxide dismutase, may also have contributed to maintenance of biomass and therefore enhanced the potential for overall coral colony survival (Thornhill et al., 2011). Therefore, the moderate increase in tissue biomass through time, despite repeat stress events, may be a result of greater resilience in corals surviving repeat bleaching events, in addition to seasonal and stress-dependent fluctuations in coral tissue (Fitt, McFarland, Warner, & Chilcoat, 2000; Wall, Ritson-Williams, Popp, & Gates, 2019).

The observed differences in antioxidant and immune activity in corals during repeat bleaching and recovery reveal that shifts in cellular priorities and mechanisms for dealing with bleaching stress exist, which ultimately shapes coral physiotypes and their trajectories through time and between stress states (Figure 5). In this case, increased catalase activity during bleaching may be linked to reactive oxygen and nitrogen species and increased host apoptosis-like pathways (Hawkins, Krueger, Becker, Fisher, & Davy, 2014; Krueger et al., 2015). Conversely, the accumulation of superoxide dismutase is decoupled from the onset and subsidence of thermal stress seen in catalase. This response may reflect the “cellular memory” of corals to thermal stress, acting as a buffer to future oxidative stress (Barshis et al., 2013; Brown et al., 2015) to reduce bleaching effects while also contributing to greater tissue retention and post-bleaching survival. Both corals and Symbiodiniaceae have a broad array of superoxide dismutase isoforms (Krueger et al., 2015; Richier, Sabourault, & Courtiade, 2006), which can be used to combat reactive oxygen species originating from damage to photomachinery and host mitochondria that together can trigger apoptosis (Dunn et al., 2012). The persistent increase of superoxide dismutase through time thus suggests antioxidant “frontloading” (Barshis et al., 2013) is an important strategy used by corals during repetitive thermal stress to reduce damage.

Engagement of the melanin pathway (collectively here as the prophenoloxidase reservoir and the melanin product) is known to be an important generalized stress response for mitigating disease and thermal stress effects in corals (Mydlarz, Holthouse, Peters, & Harvell, 2008; Palmer et al., 2010; Wall et al., 2018). Likewise, exposure to thermal stress increases antioxidant activity in corals and Symbiodiniaceae (Gardner et al., 2017). In the first-bleaching event, the melanin pathway was observed as the primary cellular response to warming in all corals, regardless of bleaching sensitivity or symbiont community. This initial peak in melanin synthesis further supports the central role of this pathway as a generalized cellular response to periodic stress, including wound repair, disease, photodamage, and bleaching, and/or a photoprotective role to reduce excess excitation energy (Mydlarz et al., 2008; Palmer, Traylor-Knowles, Willis, & Bythell, 2011; Palmer et al., 2010; Wall et al., 2018). Its subsequent decline suggests the melanin cascade may i) prime the antioxidant response and no longer is required to act as a primary stress response role, or ii) had its capacity overwhelmed in the second-bleaching, resulting in corals primarily using antioxidant stress response mechanisms. Our findings are more in line with the former. Under this thermal-priming hypothesis, sublethal thermal stress and high variable temperature regimes can contribute to protective mechanisms in the coral holobiont and bleaching resistance (Ainsworth et al., 2016; Barshis et al., 2013; Bellantuono, Granados-Cifuentes, Miller, Hoegh-Guldberg, & Rodriguez-Lanetty, 2012; Safaie et al., 2018). However, an increased frequency or magnitude of thermal stress may overcome acquired cellular responses (or symbiont partnerships) that support coral resistance to bleaching and mortality (Ainsworth et al., 2016).

In our study, the hierarchy of cellular stress responses shifts away from melanin-synthesis in favor of antioxidant activity (Figure 5), which may reflect the specialized nature of antioxidants in mitigating cellular damage and maintaining coral holobiont homeostasis (Murphy, Collier, & Richmond, 2019). Antioxidant proteins are central to cellular stress response network (Scandalios, 2002) and in corals there is clear evidence that antioxidant enzymes have a stress inducible role, driven by both environmental stress and pathogen elicitors (Mydlarz & Harvell, 2007; Mydlarz et al., 2008; Palmer et al., 2011). Indeed, *M. capitata* exposed to thermal stress showed greater antioxidant activity but lower melanin when primed by a history of high pCO_2_-variability (Wall et al., 2018). The increasing role of catalase, peroxidase, and superoxide dismutase activity through time and in the second-bleaching event, compared to attenuated melanin-synthesis pathway (Figure 5, S1, S2), further demonstrates that specialized antioxidative enzymes are important for coping with chronic or persistent stress, as often reported in thermally stressed corals (Gardner et al., 2017; Lesser, 1997). Such constitutive upregulation of cellular defenses have also been observed in corals from areas of high thermal variance (Barshis et al., 2013) and those acclimatized to warmer conditions before the onset of bleaching stress (Bellantuono et al., 2012). In particular, the continued increases in superoxide dismutase and the persistence of high levels of peroxidase in the final 2016-recovery period indicate that increasing constitutive levels of select antioxidants is key to maintaining cellular homeostasis after repeated or chronic thermal perturbations (Figure 5, S2). Such shifts in homeostatic mechanisms through strategies like gene plasticity (Kenkel & Matz, 2016) or constitutive frontloading (Barshis et al., 2013) reveal the important role of immunity, antioxidants, and their interplay in shaping ecological and evolutionary trajectories of reef corals and the future of coral reefs.

In conclusion, we show coral physiotypes were distinct during each period of bleaching and recovery from 2014-2016, and this indicates physiological legacy effects driven by chronic stress or beneficial acclimation influence both bleaching and recovery trajectories. Bleaching was strongly related to symbiont communities and their sensitivity to thermal stress, which was influenced by site environmental history. However, physiotypes were shaped by the cascade of immune and antioxidant activity in the bleaching and recovery periods irrespective of symbiont communities, providing a mechanistic hypothesis for the shifting coral physiotype baseline.

Specifically, we observed changes in constitutive immunity from one year to the next in response to extreme stress events, indicating a fundamental change in the homeostatic mechanisms employed by corals. In an era of increasing frequency and magnitude of thermal stress, the ongoing study of mechanisms such as frontloading of genes like heat shock proteins and antioxidants to attenuate bleaching effects and support thermal resilience (Barshis et al., 2013) is of great importance. Additionally, the examination of acclimatization of populations to environmental stress that may be gained through gene expression plasticity (Kenkel & Matz, 2016), or epigenetic mechanisms (Durante, Baums, Williams, Vohsen, & Kemp, 2019; Eirin-Lopez & Putnam, 2019; Liew et al., 2018) will be critical. Our study provides details of cellular and physiological legacy effects in corals and evidence that environmental memory shapes the homeostatic strategies of corals, which ultimately dictates a coral’s ability to respond to future stress events.

## Acknowledgements

The authors acknowledge funding support from an Environmental Protection Agency STAR Fellowship Assistance Agreement (FP-91779401-1) to C.B.W and NSF 1756623 (Biological Oceanography, Integrative and Ecological Physiology, and EPSCoR) to H.M.P. The views expressed in this publication have not been reviewed or endorsed by the EPA and are solely those of the authors. We also thank P.J. Edmunds for insightful comments on an earlier manuscript draft. This is SOEST contribution number xxx and HIMB contribution number xxx.

## Author contributions

C.B.W., C.A.R., L.D.M., R.D.G., and H.M.P. designed the project. C.B.W., C.A.R., L.D.M., and H.M.P. wrote the manuscript, and C.B.W. statistically analyzed the data. Coral collections were performed by C.B.W. and A.D.W. Laboratory analyses were performed by C.B.W., C.A.R., A.D.W., B.E.L., and D.E.K.

## Data availability

All data and code to generate figures and perform analyses are archived and openly available at Github (https://github.com/cbwall/Gates-Mydlarz-bleaching-recovery/releases/tag/v4).

