## Supplemental Tables and Figures for "Shifting Baselines: Physiological legacies contribute to the response of reef coral to frequent heat waves"

**Supplementary Table 1.** Statistical analysis of environmental history and bleaching event effects on *Montipora capitata* physiology.

| *Dependent variable* | *Effect* | *SS* | *df* | *F* | *P* |
|---|---|---|---|---|---|
| symbionts $cm^{-2}$ | Period | $1.108 \times 10^{13}$ | 3 | 9.470 | **<0.001** |
| | Site | $3.068 \times 10^{10}$ | 1 | 0.079 | 0.779 |
| | Symbiont | $5.640 \times 10^{13}$ | 1 | 144.647 | **<0.001** |
| | Period × Site | $4.604 \times 10^{12}$ | 3 | 3.936 | **0.009** |
| | Period × Symbiont | $3.461 \times 10^{12}$ | 3 | 2.959 | **0.033** |
| | Site × Symbiont | $8.347 \times 10^{11}$ | 1 | 2.141 | 0.144 |
| | Period × Site × Symbiont | $4.342 \times 10^{12}$ | 3 | 3.712 | **0.012** |
| | Residual | $1.174 \times 10^{14}$ | 301 | | |
| chlorophyll *a* $cm^{-2}$ | Period | 85.343 | 3 | 11.986 | **<0.001** |
| | Site | 23.851 | 1 | 10.049 | **0.002** |
| | Symbiont | 3.414 | 1 | 1.438 | 0.231 |
| | Period × Site | 7.037 | 3 | 0.988 | 0.399 |
| | Period × Symbiont | 66.062 | 3 | 9.278 | **<0.001** |
| | Site × Symbiont | 0.069 | 1 | 0.029 | 0.865 |
| | Period × Site × Symbiont | 12.746 | 3 | 1.790 | 0.149 |
| | Residual | 714.410 | 301 | | |
| chlorophyll *a* $cell^{-1}$ | Period | 0.142 | 3 | 3.310 | **0.020** |
| | Site | 0.060 | 1 | 4.189 | **0.042** |
| | Symbiont | 2.371 | 1 | 165.321 | **<0.001** |
| | Period × Site | 0.047 | 3 | 1.085 | 0.356 |
| | Period × Symbiont | 1.165 | 3 | 3.835 | **0.010** |
| | Site × Symbiont | 1.152 | 1 | 10.611 | **0.001** |
| | Period × Site × Symbiont | 0.061 | 3 | 1.429 | 0.234 |
| | Residual | 4.316 | 301 | | |
| protein $cm^{-2}$ | Period | 0.058 | 3 | 0.722 | 0.540 |
| | Site | 0.053 | 1 | 1.980 | 0.160 |
| | Symbiont | 0.126 | 1 | 4.706 | **0.031** |
| | Period × Site | 0.412 | 3 | 5.135 | **0.002** |
| | Period × Symbiont | 0.035 | 3 | 0.440 | 0.725 |
| | Site × Symbiont | 0.011 | 1 | 0.394 | 0.530 |
| | Period × Site × Symbiont | 0.048 | 3 | 0.596 | 0.618 |
| | Residual | 8.020 | 300 | | |
| total biomass $cm^{-2}$ | Period | 7.363 | 3 | 78.602 | **<0.001** |
| | Site | 0.603 | 1 | 19.312 | **<0.001** |
| | Symbiont | 0.355 | 1 | 11.359 | **<0.001** |
| | Period × Site | 1.213 | 3 | 12.954 | **<0.001** |
| | Period × Symbiont | 0.346 | 3 | 3.696 | **0.012** |
| | Site × Symbiont | 0.085 | 1 | 2.738 | 0.099 |
| | Period × Site × Symbiont | 0.065 | 3 | 0.698 | 0.554 |
| | Residual | 9.398 | 301 | | |

*Period* = Four events (first bleaching [October 2014], first recovery [February 2015], second bleaching [October 2015], second recovery [February 2016]); *Site* = two reef locations (Lilipuna and Reef 14); *Symbiont* = symbiont community dominated by *Cladocopium* sp. or *Durusdinium* sp. symbionts (C- or D-dominated). *SS* = sum of squares and *df* = degrees of freedom.

**Supplementary Table 2.** Statistical analysis of environmental history and bleaching event effects on antioxidant enzymes and immune activity of *Montipora capitata.*

| *Dependent variable* | *Effect* | *SS* | *df* | *F* | *P* |
|---|---|---|---|---|---|
| Melanin (MEL) | Period | 1.758 | 3 | 1055.892 | **<0.001** |
| | Site | $0.300 \times 10^{-3}$ | 1 | 0.556 | 0.457 |
| | Symbiont | $0.040 \times 10^{-3}$ | 1 | 0.071 | 0.790 |
| | Period × Site | 0.016 | 3 | 9.738 | **<0.001** |
| | Period × Symbiont | 0.002 | 3 | 1.311 | 0.271 |
| | Site × Symbiont | $0.017 \times 10^{-3}$ | 1 | 0.030 | 0.863 |
| | Period × Site × Symbiont | $0.297 \times 10^{-3}$ | 3 | 0.178 | 0.911 |
| | Residual | 0.165 | 298 | | |
| Prophenoloxidase (PPO) | Period | 8.112 | 3 | 207.503 | **<0.001** |
| | Site | 0.055 | 1 | 4.227 | **0.041** |
| | Symbiont | 0.054 | 1 | 4.135 | **0.043** |
| | Period × Site | 0.002 | 3 | 0.051 | 0.985 |
| | Period × Symbiont | 0.020 | 3 | 0.510 | 0.676 |
| | Site × Symbiont | $0.069 \times 10^{-3}$ | 1 | 0.005 | 0.942 |
| | Period × Site × Symbiont | 0.001 | 3 | 0.031 | 0.993 |
| | Residual | 3.857 | 296 | | |
| Peroxidase (POX) | Period | 1.657 | 3 | 16.504 | **<0.001** |
| | Site | 0.121 | 1 | 3.619 | 0.058 |
| | Symbiont | 0.001 | 1 | 0.042 | 0.838 |
| | Period × Site | 0.089 | 3 | 0.884 | 0.450 |
| | Period × Symbiont | 0.363 | 3 | 3.612 | **0.014** |
| | Site × Symbiont | 0.018 | 1 | 0.547 | 0.460 |
| | Period × Site × Symbiont | 0.014 | 3 | 0.138 | 0.937 |
| | Residual | 9.502 | 284 | | |
| Catalase (CAT) | Period | 23.590 | 3 | 118.026 | **<0.001** |
| | Site | 1.593 | 1 | 23.903 | **<0.001** |
| | Symbiont | 0.586 | 1 | 8.798 | **0.003** |
| | Period × Site | 2.096 | 3 | 10.488 | **<0.001** |
| | Period × Symbiont | 0.391 | 3 | 1.958 | 0.120 |
| | Site × Symbiont | 0.014 | 1 | 0.212 | 0.645 |
| | Period × Site × Symbiont | 0.165 | 3 | 0.862 | 0.480 |
| | Residual | 19.521 | 293 | | |
| Superoxide dismutase (SOD) | Period | $5.397 \times 10^{6}$ | 3 | 83.207 | **<0.001** |
| | Site | $0.105 \times 10^{6}$ | 1 | 4.867 | **0.028** |
| | Symbiont | $0.032 \times 10^{6}$ | 1 | 1.500 | 0.222 |
| | Period × Site | $0.038 \times 10^{6}$ | 3 | 0.578 | 0.630 |
| | Period × Symbiont | $0.093 \times 10^{8}$ | 3 | 1.430 | 0.234 |
| | Site × Symbiont | $0.008 \times 10^{6}$ | 1 | 0.378 | 0.539 |
| | Period × Site × Symbiont | $0.041 \times 10^{6}$ | 3 | 0.630 | 0.596 |
| | Residual | $6.465 \times 10^{6}$ | 299 | | |

*Period* = Four events (first bleaching [October 2014], first recovery [February 2015], second bleaching [October 2015], second recovery [February 2016]); *Site* = two reef locations (Lilipuna and Reef 14); *Symbiont* = symbiont community dominated by *Cladocopium* sp. or *Durusdinium* sp. symbionts (C- or D-dominated). *SS* = sum of squares and *df* = degrees of freedom.

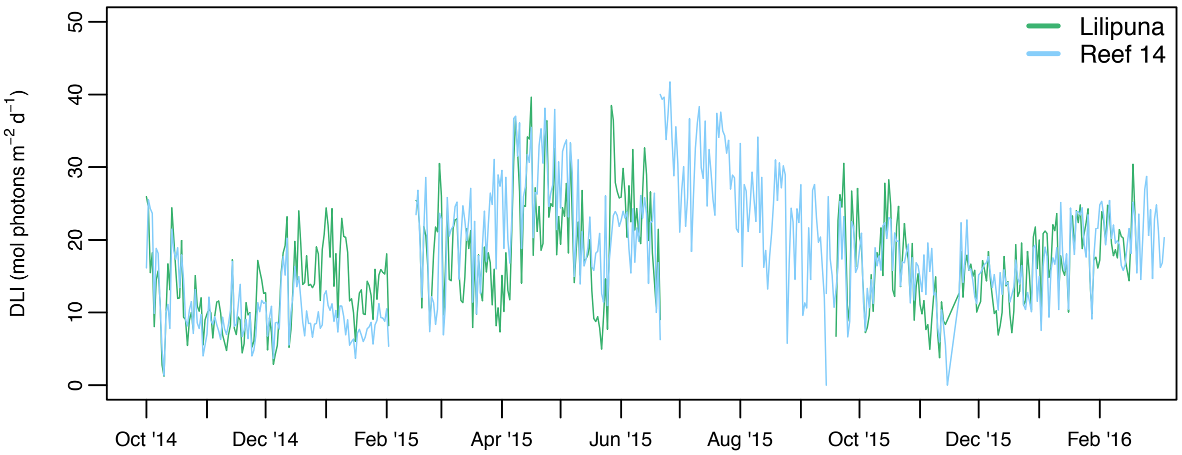


**Supplementary Figure 1.** Light availability at two reefs in Kāne‘ohe Bay during repeated bleaching and recovery periods. Photosynthetically active irradiance integrated over a 24 h day and expressed as the daily light integral (DLI) from October 2014 - March 2016. Gaps in data represent logger failure.

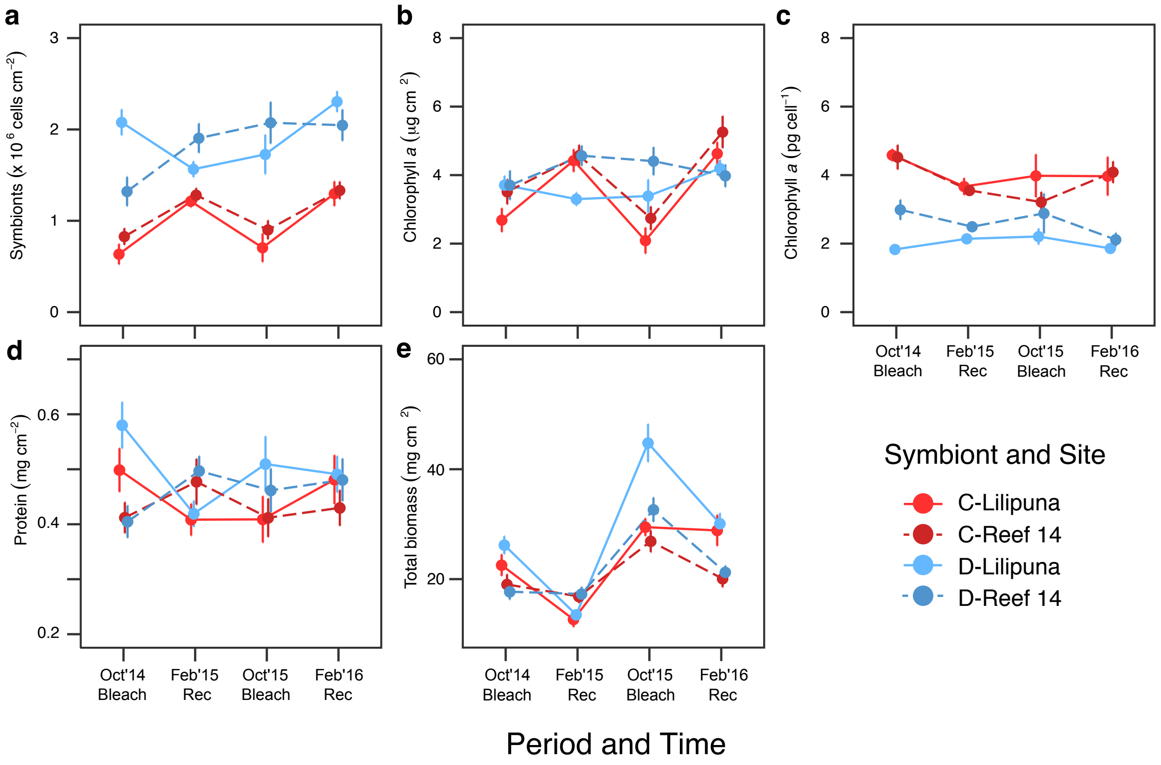


**Supplementary Figure 2.** Physiological metrics for *M. capitata* corals dominated by *Cladocopium* sp. or *Durusdinium* sp. symbionts (C or D-dominated) from two reefs in Kāne‘ohe Bay (Lilipuna, Reef 14) during repeat bleaching and recovery periods. Area-normalized (a) symbiont cell densities and (b) chlorophyll *a* concentrations (c) chlorophyll *a* per symbiont cell (d) area-normalized protein concentrations and (e) total biomass represented as ash-free dry weight. Values are mean ± SE, ($n$ = 11 – 24).

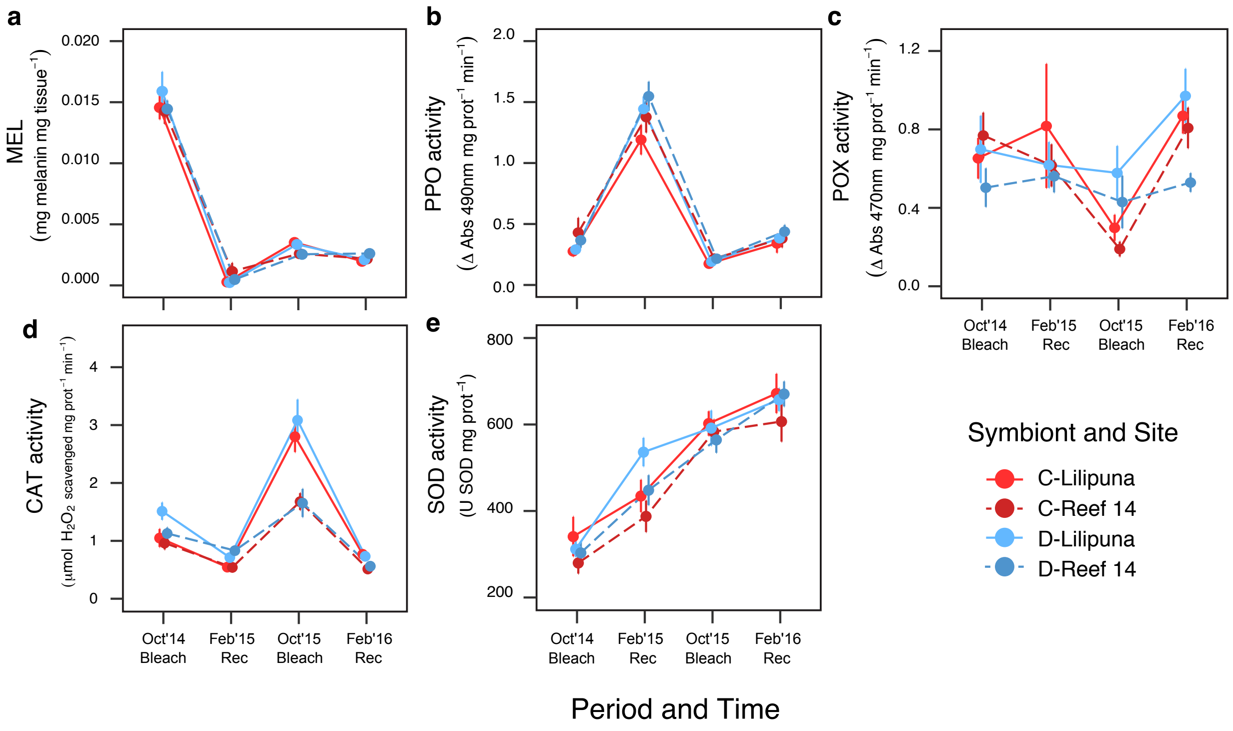


**Supplementary Figure 3.** Immunity metrics for *M. capitata* corals dominated by *Cladocopium* sp. or *Durusdinium* sp. symbionts (C or D-dominated) from two reefs in Kāneʻohe Bay (Lilipuna, Reef 14) during repeat bleaching and recovery periods. (a) Melanin (MEL), (b) prophenoloxidase (PPO), (c) peroxidase (POX), (d) catalase (CAT), and (e) superoxide dismutase (SOD). Values are mean ± SE ($n$ = 11 – 28).
